# RUNX1T1-HDAC Reprogramming of the HOX Code Signaling Drives a Targetable Pan-Cancer Lineage Plasticity

**DOI:** 10.1101/2025.10.07.681000

**Authors:** Yuyin Jiang, Siyuan Cheng, Catherine Yijia Zhang, Xiao Jin, Yaru Xu, Isaac Yi Kim, Ping Mu

**Affiliations:** Department of Urology, Yale University School of Medicine, New Haven, CT, 06511; Yale Cancer Center, Yale University School of Medicine, New Haven, CT, 06511; Department of Bioengineering, University of Illinois at Urbana-Champaign Grainger College of Engineering, Champaign, IL, 61820; Department of Population Health Sciences, Weill Cornell Medicine, Cornell University, New York, NY, 10065

**Keywords:** HOX code, Lineage plasticity, Prostate cancer, Lung cancer, AML, RUNX1T1, HDAC

## Abstract

Cancer remains a leading cause of death worldwide, with lineage plasticity emerging as a hallmark that drives therapy resistance and tumor progression by enabling cancer cells to alter identity and evade targeted therapies. Although genomic and transcriptomic aberrations correlate with lineage plasticity, the absence of pan-cancer markers to rapidly identify plastic subtypes has limited predictive utility. Homeobox (HOX) genes encode transcription factors that define tissue identity through distinct expression patterns, or HOX codes, in specific lineages. By analyzing multi-omics data, including 39 HOX genes from over 80,000 RNA-seq samples across 114 cancer types, we discovered that HOX code expression robustly represents cancer cell lineages and reveals multiple previously unrecognized lineage-plastic subtypes in prostate cancer, lung cancer, and acute myeloid leukemia (AML), each displaying altered HOX patterns compared to non-plastic subtypes. Differential analysis further identified RUNX1T1 as a novel and consistent marker of plasticity, elevated across all three cancer types and correlating with HOX code and lineage-plastic marker genes. We validated these correlations in bulk and single-cell RNA-seq from extensive preclinical and clinical cohorts and provided direct functional evidence that RUNX1T1 is required for lineage-plastic programs in prostate cancer. Using AI-based modeling, we identified NCOR/HDAC as RUNX1T1 binding partners forming a co-repressor complex that regulates HOX codes and plasticity. Finally, pharmacologic HDAC inhibition selectively suppressed the growth of plastic cells, revealing a novel therapeutic vulnerability. These findings establish ectopic RUNX1T1 as a pan-cancer biomarker and critical mediator of lineage plasticity and identify the RUNX1T1–HDAC complex as a druggable target.

## Introduction

Cancer remains the second leading cause of death worldwide, and despite advances in treatment, most cancers eventually develop drug resistance and metastasize. Cancer cells possess the ability to alter their established lineage by reverting to a stem-like state and subsequently re-differentiating into an alternative lineage, thereby evading therapies targeting their original lineage-directed survival programs [1-4]. This plasticity has been demonstrated to cause resistance to many standard cancer therapies in various types of human cancers, including prostate, breast, lung, and pancreatic cancers, as well as melanoma [2, 4-12]. Prostate cancer (PCa) represents a salient example of how cancer cells acquire resistance through lineage plasticity [13]. PCa is the second most frequently diagnosed cancer among American men, with the deadliest form being metastatic castration-resistant prostate cancer (mCRPC), which is currently incurable and is responsible for almost all PCa related death [14, 15]. Although the development of second-generation AR-targeted therapies has significantly improved the survival of men with mCRPC, resistance to these agents is unfortunately inevitable, largely limiting the clinical outcomes of patients with mCRPC [16-19]. Emerging evidence demonstrates that the luminal type of mCRPC can transform into lineage-plastic types of cancers, including neuroendocrine prostate cancer (NEPC), double-negative mCRPC, or progenitor-type mCRPC, which become independent of androgens and highly resistant to therapies. Similarly, in lung cancer, lineage plasticity contributes to the transformation of lung adenocarcinoma (LUAD) to small cell lung cancer (SCLC) or squamous cell carcinoma, resulting in resistance against EGFR inhibitors. Although less studied, lineage plasticity has been reported in other cancer types such as bladder, pancreas, esophageal cancer, and glioblastoma [20].

Efforts to decode the mechanisms driving lineage plasticity have revealed key contributors in specific cancers [21]. For example, genetic loss of *TP53* and *RB1* has been shown to facilitate lineage plasticity in prostate cancer and lung cancer [4, 22]. Across different cancer types, epigenetic regulators such as Tet Methylcytosine Dioxygenase 2 (TET2) [23] and Enhancer of Zeste 2 Polycomb Repressive Complex 2 Subunit (EZH2) [24], along with transcription factors including SRY-Box Transcription Factor 2 (SOX2) [4] and Achaete-Scute Family BHLH Transcription Factor 1 (ASCL1) [25, 26], have been reported to contribute to lineage plasticity and targeted therapy resistance in various cancers. Other epigenetic modifiers such as CHD1, LSD1, and the SWI/SNF complex [5, 7, 27-32], also contribute to the acquisition of lineage plasticity. Additionally, signaling pathways such as JAK-STAT and Notch [2, 33], RNA-binding proteins like SYNCRIP [34], and ubiquitination-related enzymes such as UBE2J1 [35] have been implicated in regulating lineage plasticity and therapy resistance in mCRPC. Despite these recent advances revealing key regulators of lineage plasticity, a pan-cancer gene marker quantifying lineage plasticity is still missing, which limits the definition and quantification of lineage plasticity across diverse cancer types and lineages. This gap in knowledge largely limits the clinical utilization of lineage plasticity as a predictive biomarker for targeted therapy resistance and hinders the practice of precision oncology.

The HOX code, defined by the spatially and temporally coordinated expression of Homeobox (HOX) genes, serves as a critical determinant of embryonic patterning and cell lineage identity during the normal developmental process [36]. In humans, the 39 HOX genes are organized into four clusters (HOXA–D), with their genomic arrangement mirroring their sequential activation during development: 3’-located genes are expressed earlier in anterior regions, while 5’-located genes are sequentially activated later in posterior regions [37]. This orchestrated expression governs anterior–posterior body segmentation and establishes tissue-specific identity, underscoring the role of the HOX code as a molecular blueprint for cellular lineage (Figure 1A) and indicating the potential link between the HOX code and lineage plasticity in PCa. Previously, we reported that lineage-plastic NEPC cells exhibit an altered HOX code compared to non-plastic PCa adenocarcinoma cells, as revealed through analysis of the Cancer Cell Line Encyclopedia (CCLE) database [38]. However, the initial exploration of the HOX code across pan-cancer types was limited due to insufficient sample size. In this study, through comprehensive bioinformatic analysis of multi-omics data, including expression profiles of 39 HOX genes from over 80,000 RNA sequencing samples across 114 cancer types, we first demonstrated that HOX codes effectively represent the lineages of cancer cells. We then identified lineage-plastic tumor subtypes in prostate cancer, lung cancer, and acute myeloid leukemia (AML), which exhibit altered HOX codes compared to non-plastic subtypes.

**Figure 1.**
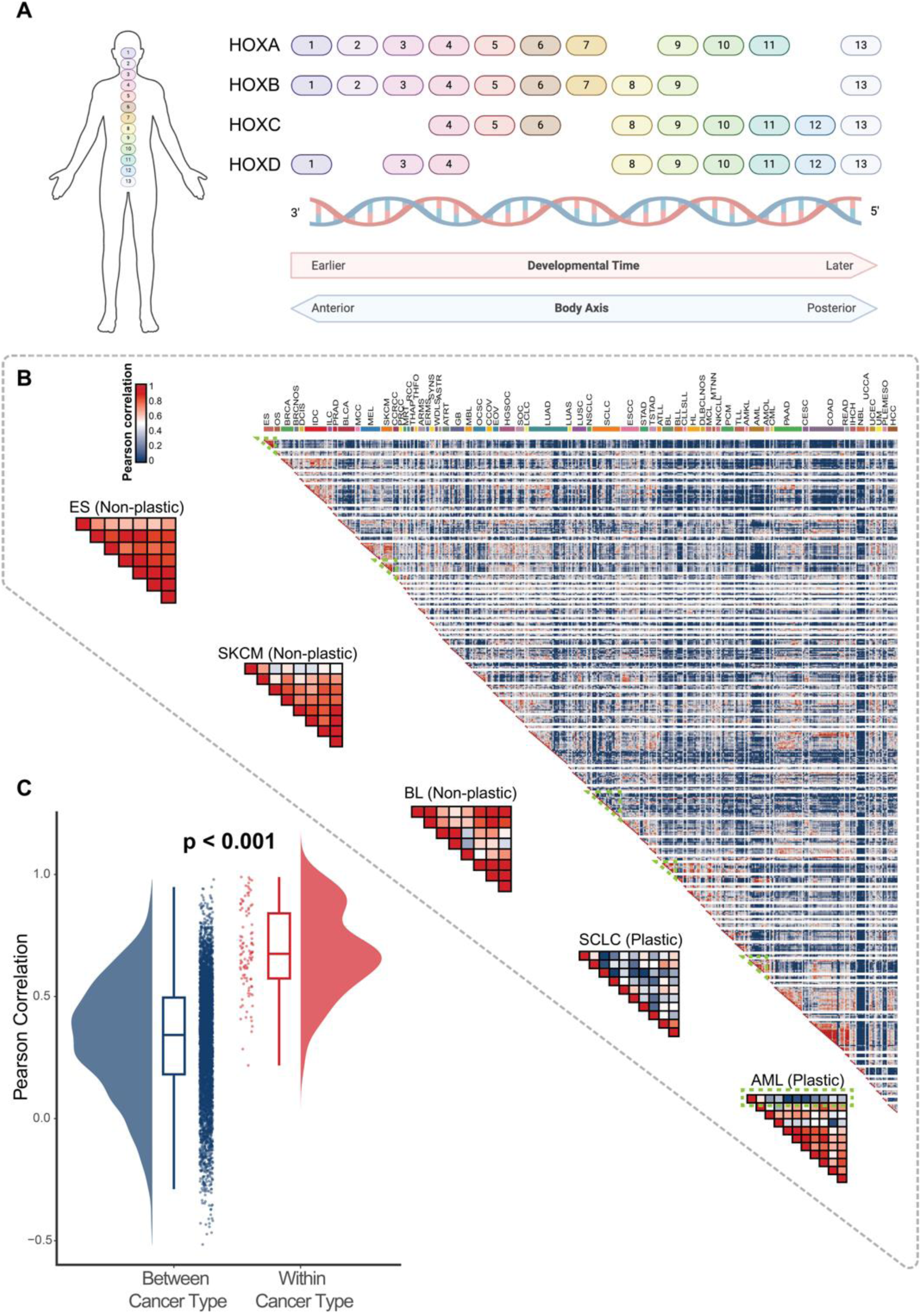
Hox code highly correlates with tissue identity and cancer cell lineages. (A) Schematic representation of HOX gene clusters (HOXA-D) arranged by embryonic developmental time and the anterior-posterior body axis. Figure was adapted from previous publication [46] and made by BioRender. (B) Heatmap showing the correlation scores between the HOX gene expression across cancer cell lines, ordered by cancer subtypes. Specific portions of the heatmap representing Ewing sarcoma (ES), cutaneous melanoma (SKCM), and Burkitt lymphoma (BL), small cell lung cancer (SCLC), and acute myeloid leukemia (AML) are outlined in green and zoomed in for better visualization and comparison. (C) Violin and box plots comparing the distribution of Pearson correlation scores between cell lines from the same cancer subtype ("Within") versus those from different cancer subtypes ("Between"). Correlations are significantly higher within the same cancer subtype (*p* < 0.001, Wilcoxon rank-sum test).

Strikingly, we identified RUNX1T1 as a novel and consistent marker of lineage plasticity, significantly elevated across all three cancer types and highly correlated with both HOX code expression and lineage-plastic transcriptomic programs. Importantly, we not only validated these correlations in bulk and single-cell RNA-seq from extensive preclinical and clinical cohorts, but also provided direct functional evidence that RUNX1T1 is required for lineage-plastic programs in prostate cancer cell lines. Using AI-based modeling, we further identified NCOR/HDAC as RUNX1T1 binding partners, forming a co-repressor complex that regulates HOX codes and lineage plasticity. Finally, we demonstrated that pharmacologic HDAC inhibition selectively suppresses the growth of plastic cells, establishing a novel therapeutic vulnerability. Together, our findings establish ectopic RUNX1T1 as a pan-cancer biomarker and critical mediator of lineage plasticity, identify the RUNX1T1–HDAC complex as a druggable target, and introduce the HOX code as a robust framework for characterizing lineage plasticity across cancers. These insights fill a major gap in leveraging lineage plasticity as a predictive biomarker for therapy resistance and provide a new strategy for therapeutically targeting these aggressive cancer subtypes with lineage plasticity.

## Results

### HOX Codes Represent Cancer Lineage Identity

In developmental biology, it is well established that the HOX code can represent cell lineage and infer the cell fate [36]. To study whether HOX code can also represent cancer cell lineage, we first accessed our previously established pan-cancer cell line database, PCTA, which integrates high-quality transcriptomic data from over 80,000 RNA-seq samples across 535 cancer cell lines [39, 40]. Using these RNA-seq samples, we generated HOX codes unique to each cancer cell line, defined as the median expression of 39 HOX genes from all RNA-seq samples of the same cancer cell line (Table 1). Based on established scientific evidence, these cancer cell lines can be categorized into specific cancer subtypes (e.g., prostate adenocarcinoma, PRAD) and further classified by their anatomic site of origin (e.g., prostate). To evaluate the conservation of HOX codes within and across cancer subtypes, we performed pairwise Pearson correlation analyses and visualized the results as a heatmap, ordering cell lines by cancer subtype. As we expected, most cancer subtypes exhibited robust intra-cancer subtype HOX code conservation, with Ewing sarcoma (ES), cutaneous melanoma (SKCM), and Burkitt lymphoma (BL) displaying particularly strong coherence (Figure 1B). For a more quantitative assessment, we compared HOX code correlations between cell lines from the same cancer subtype versus those from different cancer subtypes. Correlations were significantly higher within the same cancer subtype than between different subtypes (*p* = 0.002, one-sided Wilcoxon rank-sum test; Figure 1C). These findings suggest that HOX codes may serve as markers of cancer lineage identity, extending their role from developmental biology to oncology.

**Table 1.**
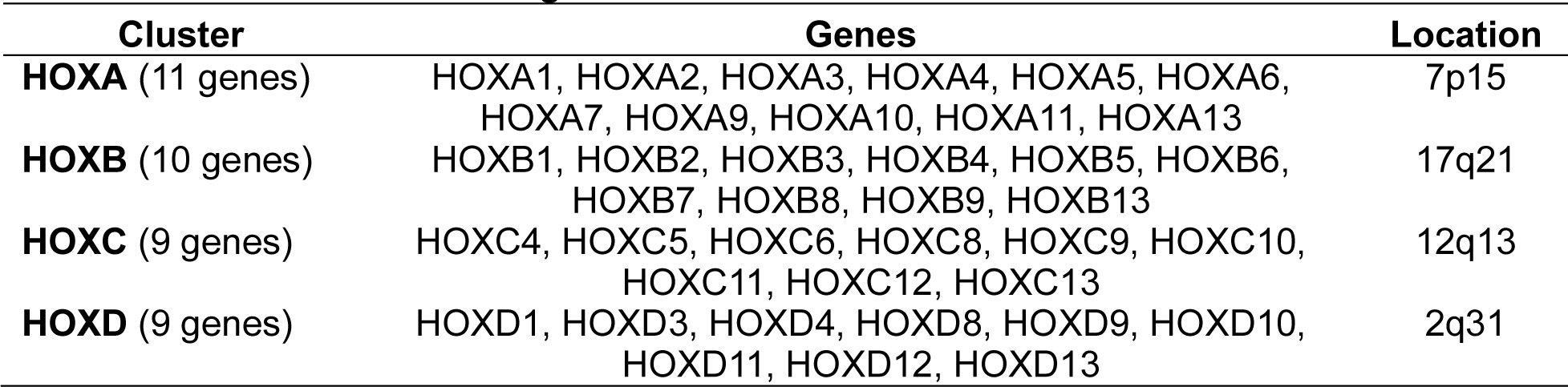
List of 39 human HOX genes.

### HOX Codes Identified Lineage-Plastic Subtypes in Prostate Cancer, Lung Cancer, and AML

Most precision medicine strategies target cancer cells based on lineage-defining markers, such as AR in prostate cancer, EGFR in lung cancer, and ER in breast cancer. However, these approaches frequently fail because cancer cells can escape targeted therapy by adopting alternative lineages, a phenomenon known as lineage plasticity, which is increasingly reported in the clinic [2, 4-10]. A major reason we cannot systematically capture this phenotype is that “cancer lineage” is still defined almost exclusively by subtype-specific markers. Such markers describe the presence of one lineage state, but they provide no framework for recognizing when a tumor has switched to another lineage. To overcome this limitation, we investigated whether HOX codes could provide a pan-cancer framework for identifying lineage-plastic subtypes in different cancers. We first defined lineage plastic cancer cell lines as a marked divergence in HOX code relative to other cell lines from the same cancer subtype or anatomic origin, quantified by reduced Pearson correlation of median HOX gene expression. This analysis revealed two distinct patterns of lineage plasticity. In acute myeloid leukemia (AML), lineage-plastic subtypes were rare, with most AML cell lines exhibiting highly similar HOX codes, consistent with conserved lineage identity (Figure 1B). In contrast, small cell lung cancer (SCLC) displayed pervasive HOX code heterogeneity, with most cell lines showing low pairwise correlations and divergent lineage-associated HOX expression profiles (Figure 1B). These observations are supported by within-group correlation distributions and hierarchical clustering based on HOX code similarity at both the cancer-type and sample-level resolution.

However, we recognize that some cancer subtypes may arise from the same anatomical site, a detail not fully captured in our initial analysis. To address this, we extended our investigation by analyzing HOX codes across different sites of cancer origin. Specifically, we generated HOX codes specific to each cancer subtype and evaluated correlation between cancer subtypes by comparing their unique HOX codes. As expected, this analysis revealed that most anatomic sites harbored cancer subtypes with high intra-site HOX code correlations, likely due to the shared developmental origins of those cancers (Figure 2A). In contrast, cancer subtypes arising from different organs exhibited low inter-site HOX code correlations (*p* = 0.002, one-sided Wilcoxon rank-sum test), suggesting distinct lineage origins or potential lineage plasticity. For example, breast neoplasms (BNNOS) and kidney malignant rhabdoid tumors (MRT) showed markedly different HOX patterns because they develop from distinct precursor cells [41, 42]. Applying our previous definition of lineage plasticity, we were able to identify a third lineage-plastic pattern exhibited by prostate cancer. Specifically, two types of prostate cancers, prostate adenocarcinomas (PRAD) and prostate small cell carcinomas (PRSCC), displayed markedly different HOX codes even though they both originated in the prostate. This difference likely occurs because PRSCC develops from PRAD through a process called trans-differentiation, where cancer cells switch their identity (lineage plasticity) [4]. To examine HOX expression changes associated with plasticity, we visualized gene-level expression patterns across representative cell lines in prostate cancer, lung cancer, and AML (Figure 2B). Notably, in prostate cancer, the lineage-plastic H660 cell line showed increased expression of anterior HOX genes (e.g., HOXB2, HOXB3) and decreased expression of posterior HOX genes (e.g., HOXB13) relative to non-plastic lines. A similar anterior-to-posterior shift in HOX expression was observed in AML. In lung cancer, although variability was higher, lineage-plastic cell lines also showed a trend of reduced anterior and elevated posterior HOX gene expression compared to their non-plastic counterparts (Figure 2B). Collectively, these findings suggest that HOX codes could serve as markers that represent lineage identity, while changes in these markers indicate when cancer cells switch identities and become lineage plastic.

**Figure 2.**
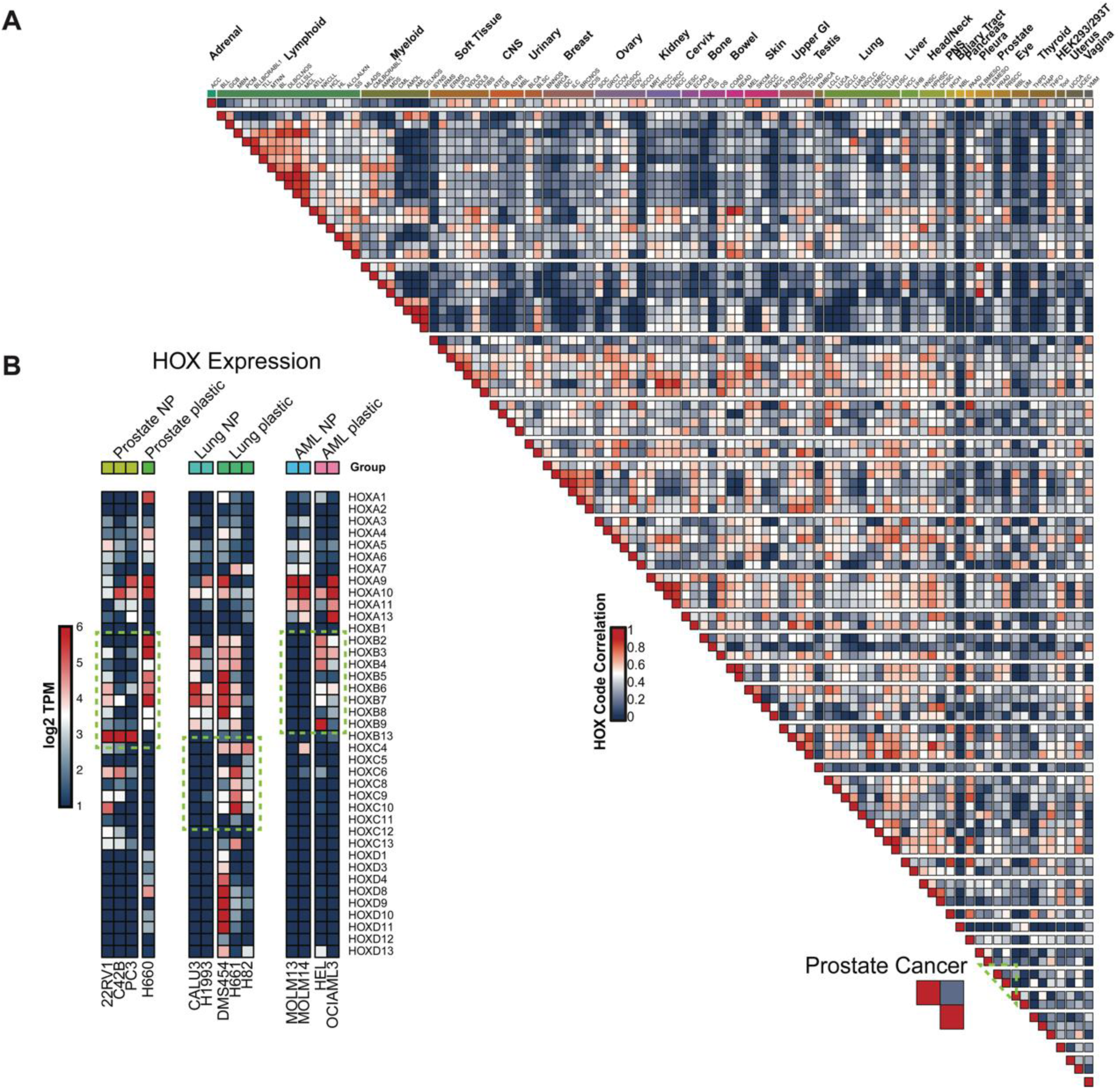
Variations in HOX gene expression patterns are highly relevant biomarkers of lineage-plastic cancers. (A) Heatmap showing the correlation scores between the HOX gene expression across cancer subtypes, ordered by anatomic sites. Heatmap representing prostate cancer is outlined in green and zoomed in. (B) Heatmap of HOX gene expression across prostate cancer, lung cancer, and AML cell lines separated in three clusters. The green box highlights distinct HOX gene expression patterns in lineage-plastic vs. non-plastic cell lines. NP (non-plastic)

### HOX Codes Identify Preclinical Models for Dissecting Pan-Cancer Lineage Plasticity

To systematically identify lineage-plastic cancer subtypes on a pan-cancer scale, we developed a global analysis to visualize the conservation of HOX code gene expression patterns. We reasoned that cancer subtypes with a single, stable lineage would exhibit a unimodal distribution of intra-site HOX code gene expression correlations. In contrast, cancer subtypes containing multiple lineage distinct subclones or undergoing active lineage transition would display a multimodal distribution, indicating significant lineage heterogeneity. To visualize this, we employed a circular “Kiwi” plot that displays the pairwise correlation distributions for cancer subtypes grouped by their anatomic site of origin. Each segment shows the distribution of correlations as a half-violin plot, allowing for the detection of multiple peaks (local maxima) that signify the presence of distinct lineage states within a single anatomic site. Cancer subtypes with various lineage are ordered by the number of detected peaks, providing a quantitative ranking of their heterogeneity in lineage. This visualization clearly distinguished cancers with a single, highly conserved HOX code expression pattern distribution (unimodal) from those displaying substantial heterogeneity (multimodal). Importantly, this approach confirmed that cancers known to contain lineage-plastic subtypes, such prostate, lung, and myeloid cancers, exhibit multimodal distributions, indicating pronounced HOX code divergence and the presence of multiple lineage states (Figure 3A).

**Figure 3.**
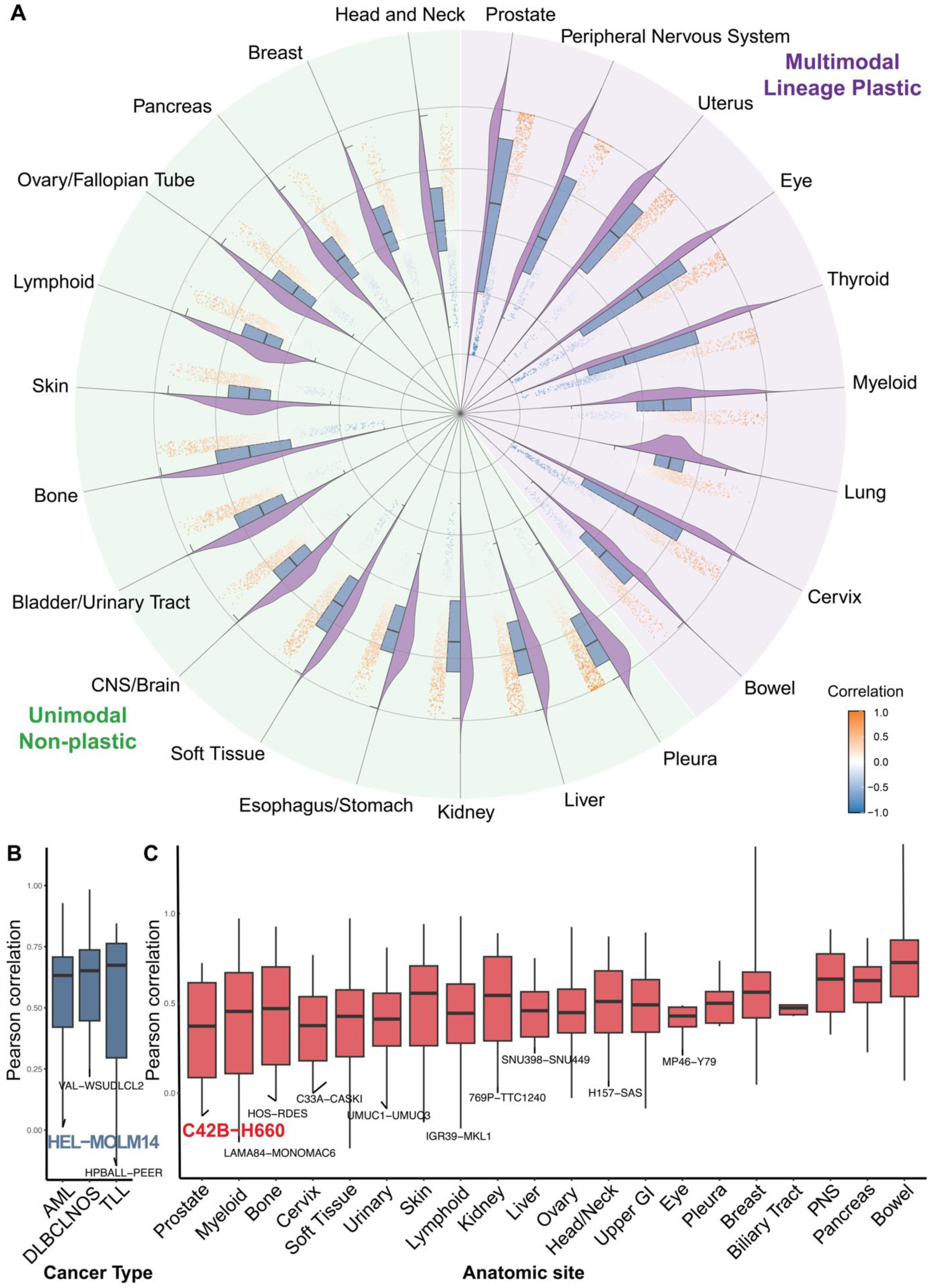
HOX codes identify lineage-plastic and molecular subtypes in prostate cancer, lung cancer, and AML. (A) “Kiwi” plot (circular comparison plot) providing a global overview of pairwise HOX gene correlation distributions across cancer subtypes of various lineages. Each segment represents a distinct lineage, with the left half–violin plot (purple) showing the distribution of correlations and the right half–boxplot (blue) indicating the median and interquartile range. Cancer subtypes with a single, stable lineage exhibit a unimodal distribution of intra-site HOX code gene expression correlations, whereas subtypes containing multiple lineage-distinct subclones or undergoing active lineage transitions display multimodal distributions, indicative of significant lineage heterogeneity. Peak counts were estimated by local maxima detection, and lineage subtypes are ordered by decreasing number of peaks. Statistical significance was assessed using a two-sided Wilcoxon rank-sum test (p < 0.05, ***). (B) Comparison of intra-cancer HOX code expression correlation scores across selected cancer subtypes. The low correlation between the HEL and MOLM14 acute myeloid leukemia (AML) cell lines highlights their high degree of lineage heterogeneity. (C) Comparison of intra-cancer HOX code expression correlation scores across selected prostate cancer and lung cancer subtypes and anatomic sites. The low correlation between the C4-2B and H660 prostate cancer cell lines illustrates pronounced lineage heterogeneity.

To further dissect lineage-plastic subtypes in prostate cancer, lung cancer, and acute myeloid leukemia (AML), we visualized HOX code similarity among representative cancer cell lines using Pearson correlation, highlighting pairs with the lowest correlations of HOX code expression patterns within each cancer subtype and organ site (Figure 3B and 3C). As expected, in AML, the correlation of HOX code expression between the lineage-plastic HEL cell line and the non-plastic MOLM14 cell line was the lowest among all AML cell line pairs (Figure 3B), indicating marked divergence in lineage identity. Similarly, in prostate cancer, the correlation of HOX code expression between the lineage-plastic H660 cell line and the non-plastic C4-2B cell line showed the greatest dissimilarity among prostate-derived models (Figure 3C). These findings highlight HEL and H660 as representative lineage-plastic models, with HOX code divergence serving as a marker of plasticity. Collectively, these analyses demonstrate that HOX code divergence robustly captures lineage-plasticity across multiple cancer types and model systems.

### Elevated RUNX1T1 Expression is Correlated with Lineage-Plastic Subtypes Compared to Their Non-Plastic Counterparts

Our analysis of HOX code patterns revealed that prostate cancer, lung cancer, and acute myeloid leukemia (AML) exhibit clear evidence of lineage plasticity. To identify genes associated with the acquisition of lineage plasticity across these cancer subtypes, we retrieved all samples derived from the relevant cell lines and performed differential gene expression analysis between lineage-plastic and non-plastic cancer subtypes and cell lines. This analysis revealed distinct transcriptional signatures associated with plasticity (log2 fold change > 3 and adjusted p < 0.01; Figure 4A–C). A Venn diagram of these results identified shared gene expression patterns across the three cancer subtypes, with 269 genes significantly upregulated in lineage-plastic prostate cancer, lung cancer, and AML (Figure 4D). Notably, RUNX1T1 emerged as a top differentially expressed gene, consistently and significantly upregulated in all three lineage-plastic cancer subtypes (Figure 4E).

**Figure 4.**
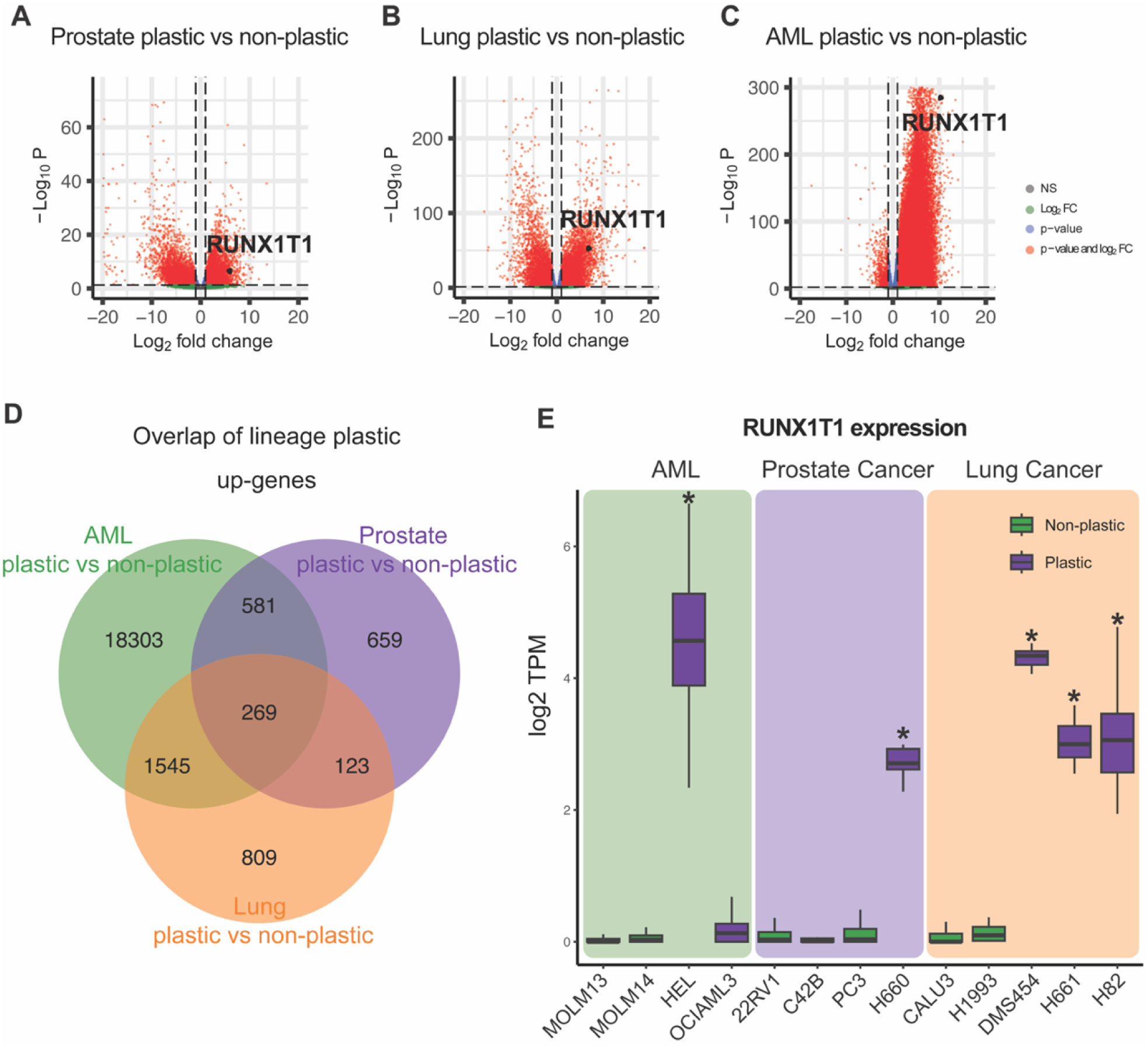
RUNX1T1 expression is highly elevated in lineage-plastic cell lines compared to their non-plastic counterparts. (A-C) Volcano plots comparing gene expression profiles between lineage-plastic and non-plastic subtypes within prostate cancer, lung cancer, and AML. A total of 53,175 genes were analyzed. RUNX1T1 is significantly upregulated in all lineage-plastic subtypes and cell lines across the three cancer types. (D) Venn diagram showing 269 significantly upregulated genes shared among plastic prostate cancer, lung cancer, and AML. (E) Bar plot comparing RUNX1T1 expression levels across lineage-plastic and non-plastic cell lines. Asterisks indicate significantly higher expression in plastic cell lines within each cancer subtype.

To further investigate the association between RUNX1T1 expression and the lineage-plastic phenotype, we expanded our analyses to include additional models. Using qRT-PCR in prostate cancer cell lines, we confirmed that lineage-plastic cell lines (H660, LASCPC1) displayed significantly elevated levels of RUNX1T1, concurrent with high expression of anterior HOXB genes and loss of the posterior HOXB13 (Figure 5A). Moreover, we validated that elevated RUNX1T1 expression was not only associated with altered HOX code expression patterns but also with canonical prostate lineage-plasticity markers, including downregulation of androgen receptor (AR) signaling and upregulation of stem-like, epithelial-to-mesenchymal transition (EMT), and neuroendocrine (NE) marker expression (Figure 5B). In addition, we assessed protein expression and found that, consistent with the RNA-level data, RUNX1T1 protein was elevated in lineage-plastic cancer cells from both prostate and lung cancers (Figure 5D).

**Figure 5.**
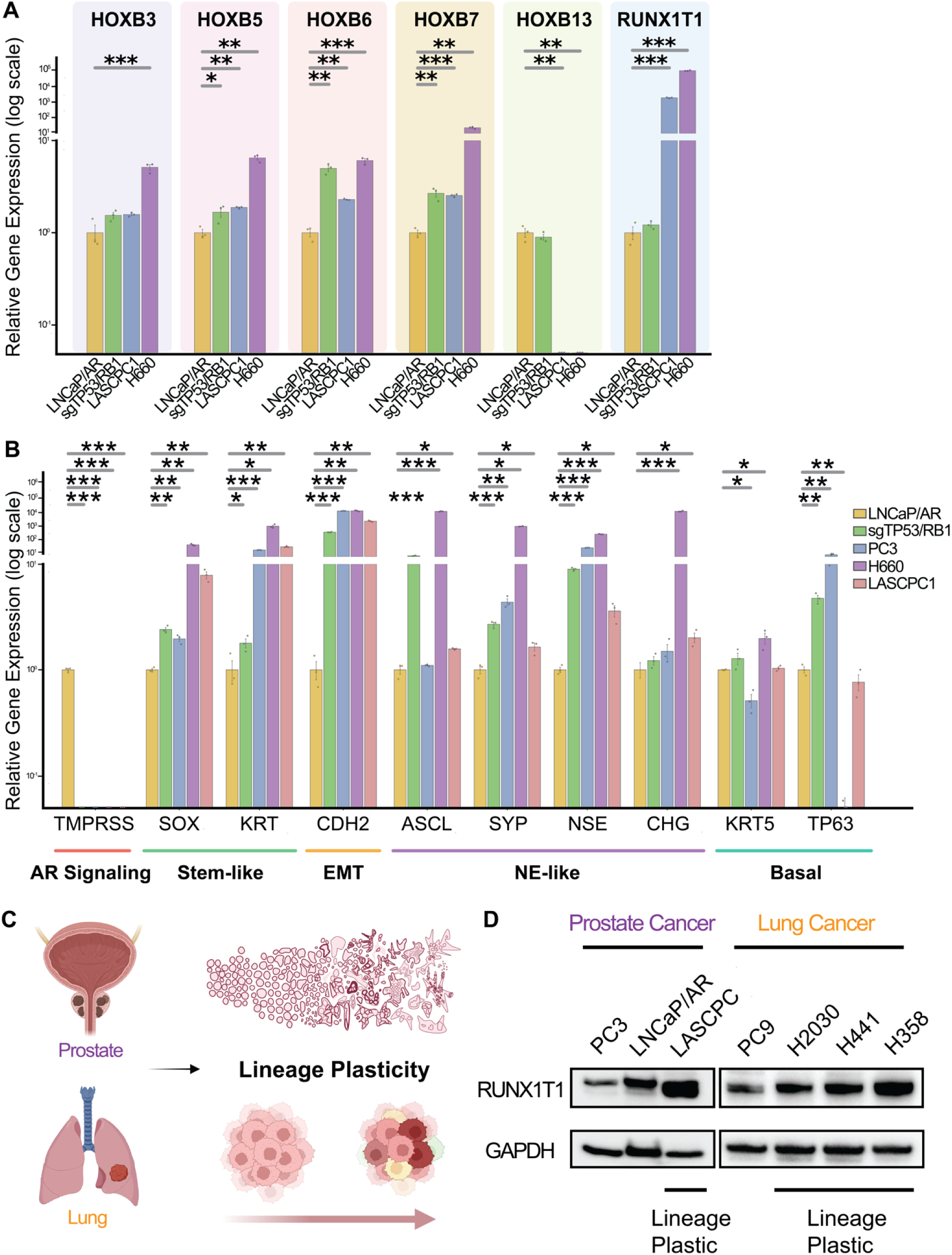
Correlated upregulation of RUNX1T1 and HOX code expression is validated in lineage-plastic cancer subtypes compared to their non-plastic counterparts. (A) qRT-PCR analysis of HOX and RUNX1T1 genes expression in a panel of prostate cancer cell lines. Lineage-plastic lines (H660, LASCPC1) show significantly elevated anterior HOXB genes (such as HOXB3) with a concomitant loss of the posterior gene HOXB13. The result also showed RUNX1T1 expression was elevated in lineage plastic cells compared to their non-plastic counterparts. (B) qRT-PCR analysis of key lineage marker genes, showing downregulation of AR signaling genes and upregulation of lineage plastic markers including stem-like, EMT, neuroendocrine, and basal epithelial markers in plastic cell lines. (C) Schematic illustrating the concept of lineage plasticity, where cancer cells from certain lineage with specific histology organization can trans-differentiate into a lineage-plastic state. (D) Western blot results demonstrating higher RUNX1T1 protein levels in lineage-plastic prostate (LASCPC1) and lung (H2030, H441, H358) cancer cell lines compared to their non-plastic counterparts.

Building on the association of RUNX1T1 with lineage plasticity, we next examined its expression in human patient samples and mouse model–derived single-cell RNA sequencing (scRNA-seq) data to determine whether the bulk-level findings were recapitulated at the single-cell level. Dimensional reduction plots revealed distinct RUNX1T1 expression patterns between lineage-plastic and non-plastic cancers (Figure 6A and 6E). Specifically, lineage-plastic neuroendocrine prostate cancers showed higher RUNX1T1 expression, whereas non-plastic adenocarcinomas exhibited lower RUNX1T1 expression in both human and mouse models (Figure 6B, 6D, 6F, and 6G). Statistical comparisons using Wilcoxon rank-sum tests confirmed significant differences in RUNX1T1 expression between plastic and non-plastic groups in both species (Figure 6D and 6G, p < 0.05). Consistently, UMAP analysis of HOXB gene expression in the human single-cell model corroborated the bulk-level HOX code patterns: the non-plastic PRAD cluster showed high expression of HOXB13, a marker of prostatic identity, whereas the plastic PRSCC cluster lost HOXB13 expression and instead upregulated HOXB3 and HOXB5 (Figure 6C). Together, these results suggest that elevated RUNX1T1 expression represents a robust and consistent signature of lineage plasticity across multiple cancer cell lines as well as in both mouse and human single-cell datasets.

**Figure 6.**
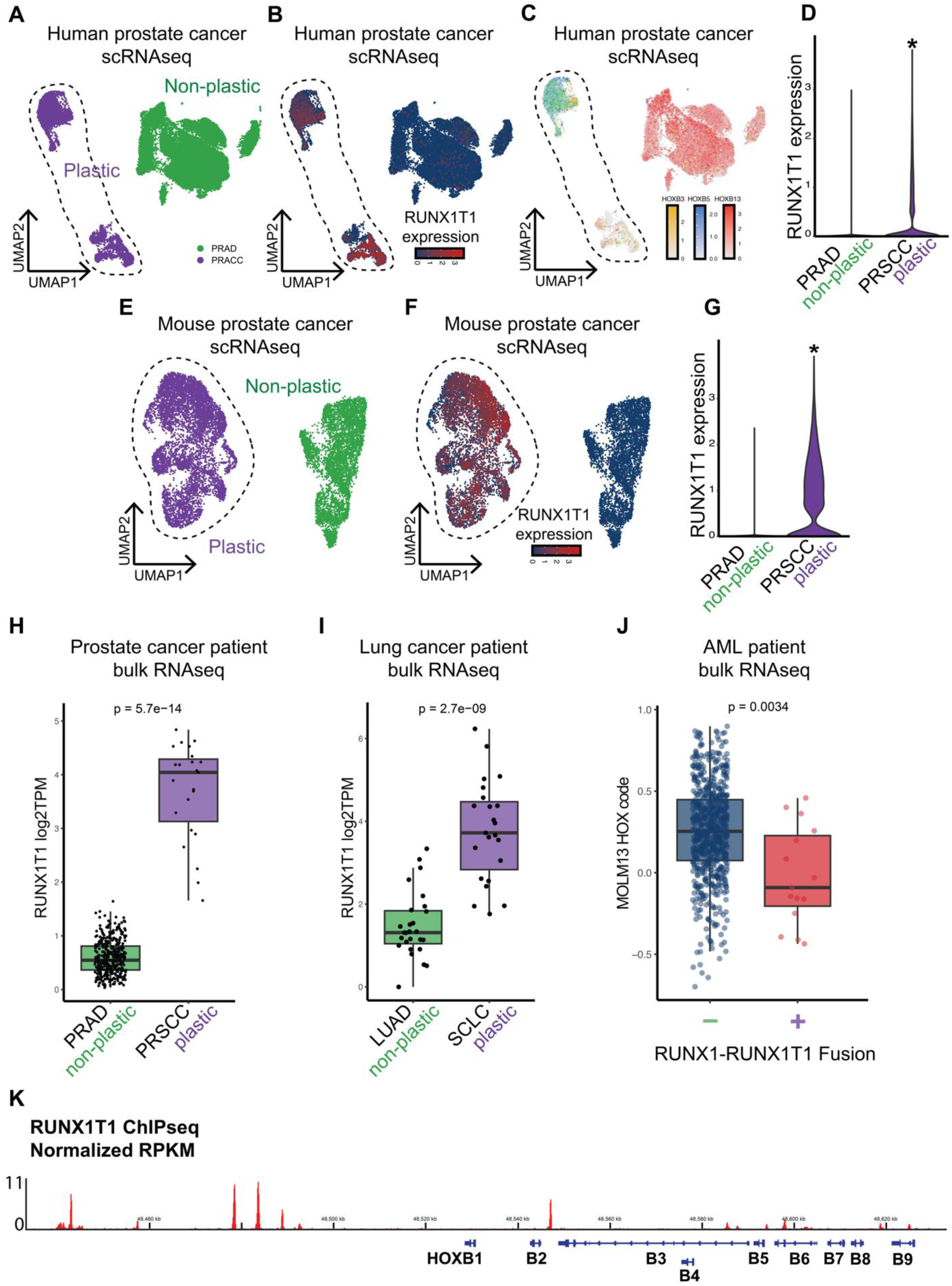
Expression of RUNX1T1 is highly upregulated in lineage plastic cancer subtypes compared to the non-plastic counterparts in both mouse and patient samples. (A) UMAP plots from single-cell RNA sequencing (scRNA-seq) analysis of human single-cell models. Green cluster represents non-plastic adenocarcinoma prostate cancer and purple cluster represents plastic neuroendocrine prostate cancer. PRAD:prostate adenocarcinomas; PRSCC:prostate small cell carcinomas. (B) UMAP plot of RUNX1T1 expression of human single-cell model showing higher RUNX1T1 expression in lineage plastic cluster. (C) UMAP analysis of HOXB gene expression of human single-cell model showing loss of HOXB13 and upregulation of HOXB3 and HOXB5 in lineage plastic prostate cancer. (D) Violin plots highlighting greater RUNX1T1 expression and variability in PRSCC of human single-cell model. (E) UMAP plots from scRNA-seq analysis of mouse single-cell models. (F) UMAP plot of RUNX1T1 expression of mouse single-cell model. (G) Violin plots of RUNX1T1 expression in mouse PRAD and PRSCC. (H) Comparison of RUNX1T1 expression between lineage plastic PRSCC and non-plastic PRAD patient samples in prostate cancers. (I) Comparison of RUNX1T1 expression between lineage plastic SCLC and non-plastic LUAD patient samples in lung cancers. (J) Pearson correlation analysis of RUNX1-RUNX1T1-fused (+) and non-RUNX1-RUNX1T1-fused (-) AML samples against the non-plastic MOLM13 HOX code, showing reduced correlation in RUNX1T1-fused samples. (K) ChIP-sequencing result of the RUNX1T1 protein.

A similar trend was observed in bulk RNA-seq data from patient samples of prostate cancer, lung cancer, and AML. We compared RUNX1T1 expression between lineage-plastic and non-plastic patient samples in prostate and lung cancers (Figure 6H and 6I). In both cancer types, the lineage-plastic subtypes, PRSCC in prostate cancer and SCLC in lung cancer, exhibited significantly higher RUNX1T1 expression compared to their non-plastic counterparts (PRAD and LUAD, respectively), as assessed using two-sided Wilcoxon rank-sum tests on log-transformed TPM values (p < 0.05). In AML, interpretation of RUNX1T1 expression was complicated by the presence of the RUNX1–RUNX1T1 fusion, which resulted in RUNX1T1 expression data being unavailable in most of the cohorts analyzed. To address this limitation, we separated patient samples into two groups, those with the fusion and those without, and compared their HOX code similarity to the non-plastic MOLM13 reference cell line. This analysis tested whether the association between RUNX1T1 and plasticity persisted despite the confounding effects of the fusion. As expected, patient samples without the fusion showed higher correlation with the non-plastic HOX code, whereas those with the fusion exhibited significantly lower correlation (Figure 6J), as determined by Wilcoxon rank-sum test (p < 0.01), indicating a transition of non-plastic to plastic characteristics due to RUNX1T1 upregulation. This finding aligns with our observations in prostate and lung cancers, reinforcing RUNX1T1 as a pan-cancer marker of lineage plasticity across multiple cancer subtypes. Finally, visualization of publicly available RUNX1T1 ChIP-seq data identified binding of the RUNX1T1 protein at the human HOXB genomic locus (Figure 6K), suggesting a direct regulatory role of RUNX1T1 in the HOX gene expression changes observed throughout this study.

### RUNX1T1 Knockdown Reverses the Changes of HOX Gene Expression and Lineage Plasticity Phenotypes

Having established a strong correlation between RUNX1T1 and lineage plasticity, we next performed functional experiments to determine if RUNX1T1 is required for maintaining the lineage plastic state. We first used siRNA to knock down RUNX1T1 in a TP53/RB1 co-deficient prostate cancer cell line, an established model genetically engineered to induce the onset of lineage plasticity [4, 22]. In this model, RUNX1T1 knockdown led to a broad reversal of lineage plasticity markers including the decreased expression of prostate basal epithelial markers and increased expression of AR-pathway and prostate luminal epithelial genes (Figure 7A) and a significant alteration of the HOX gene signature (Figure 7B).

**Figure 7.**
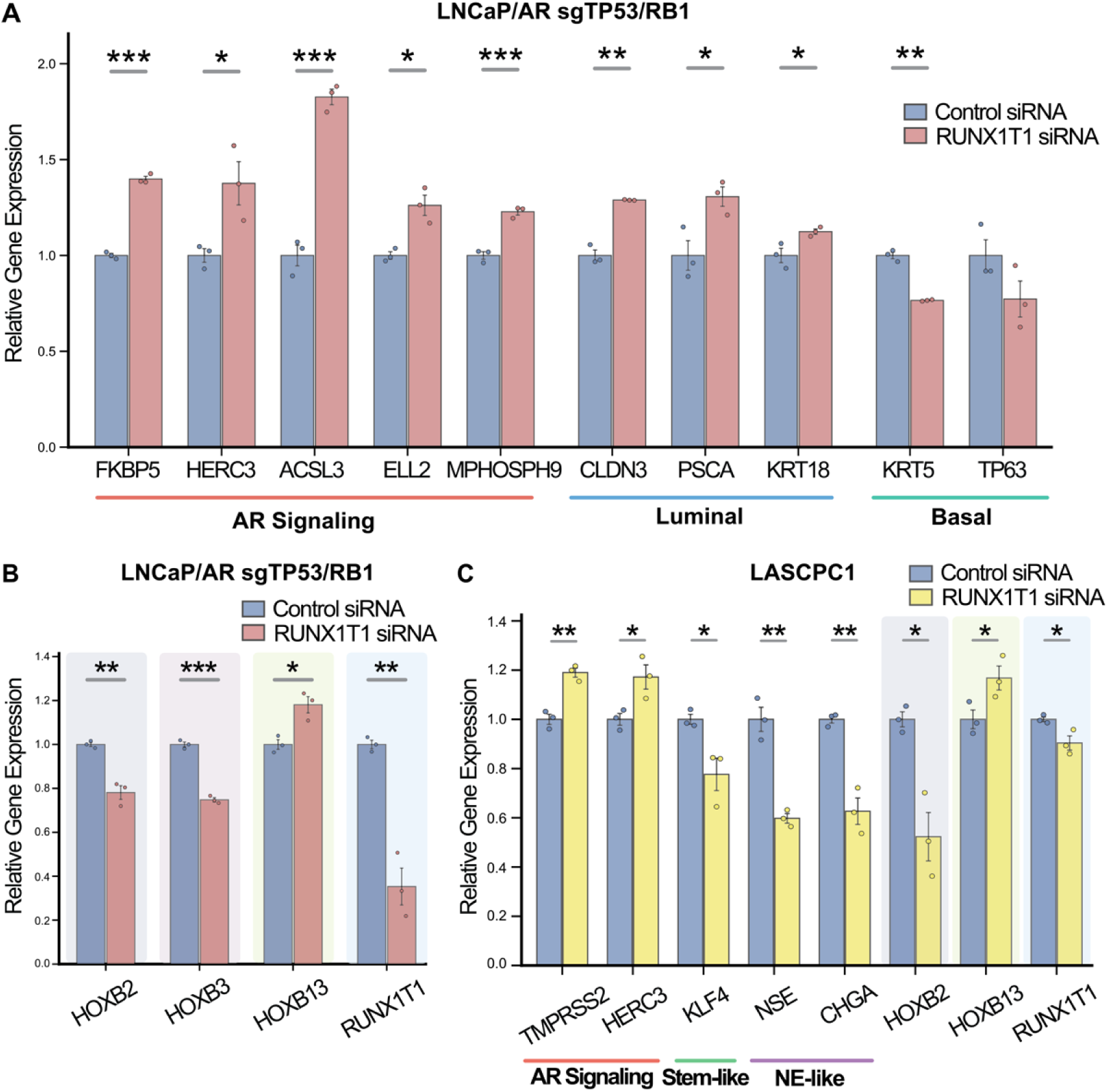
RUNX1T1 is required for altered HOX gene expression and the lineage-plastic phenotype. (A) qRT-PCR analysis in the sgTP53/RB1 prostate cancer cell line after siRNA-mediated RUNX1T1 knockdown, showing altered expression of canonical lineage marker genes. Results demonstrate restoration of AR signaling and luminal lineage genes, along with downregulation of basal lineage genes. (B) qRT-PCR analysis in the sgTP53/RB1 prostate cancer cell line after siRNA-mediated RUNX1T1 knockdown, showing altered expression of key HOX code genes. Results demonstrate downregulation of HOXB2 and HOXB3, accompanied by upregulation of HOXB13, representing a reversal of HOX code gene expression observed in TP53/RB1-DKO cells. (C) qRT-PCR analysis in the lineage-plastic LASCPC1 prostate cancer cell line following siRNA-mediated RUNX1T1 knockdown. Knockdown of RUNX1T1 reversed the elevated expression of stem-like and NE-like markers and restored HOX gene expression patterns in the lineage-plastic LASCPC1 cell line.

To confirm these findings in another model, we performed siRNA knockdown in the LASCPC1 cell line, a fully developed lineage-plastic prostate cancer model. We observed only a modest reduction of RUNX1T1, which is a known technical challenge in suspension cell lines and may also reflect strong selective pressure against its complete loss in a model fully dependent on the lineage-plastic state. Despite this partial reduction in gene expression level, we observed a significant biological effect of RUNX1T1 knockdown. This included a marked decrease in the expression of stem-like and neuroendocrine markers and a partial reversal of the characteristic HOX code, with downregulation of the anterior HOXB2 gene and upregulation of the posterior HOXB13 gene (Figure 7C). Together, these results demonstrate that RUNX1T1 expression is critical and functionally required for sustaining cancer lineage plasticity.

### RUNX1T1 Interacts with the NCOR/HDAC Complex to Regulate the Epigenetic State and Confer a Therapeutic Vulnerability

To elucidate the mechanism by which RUNX1T1 mediates lineage plasticity, we investigated its potential protein–protein interactions. Using the AI-based model AlphaFold-3, we performed interactome prediction based on the structures of RUNX1T1 in complex with known epigenetic modifiers [43]. These AI-based structural predictions suggested, with high confidence, that RUNX1T1 forms a tripartite complex with the nuclear receptor corepressors NCOR1/2 and histone deacetylases HDAC1–3 (Figure 8A). Importantly, the predicted Template Modeling (pTM) and interface predicted Template Modeling (ipTM) scores confirmed a high-confidence interaction interface for the RUNX1T1–NCOR1/2–HDAC complex, substantially stronger than those predicted for bipartite interactions (Figure 8B).

**Figure 8.**
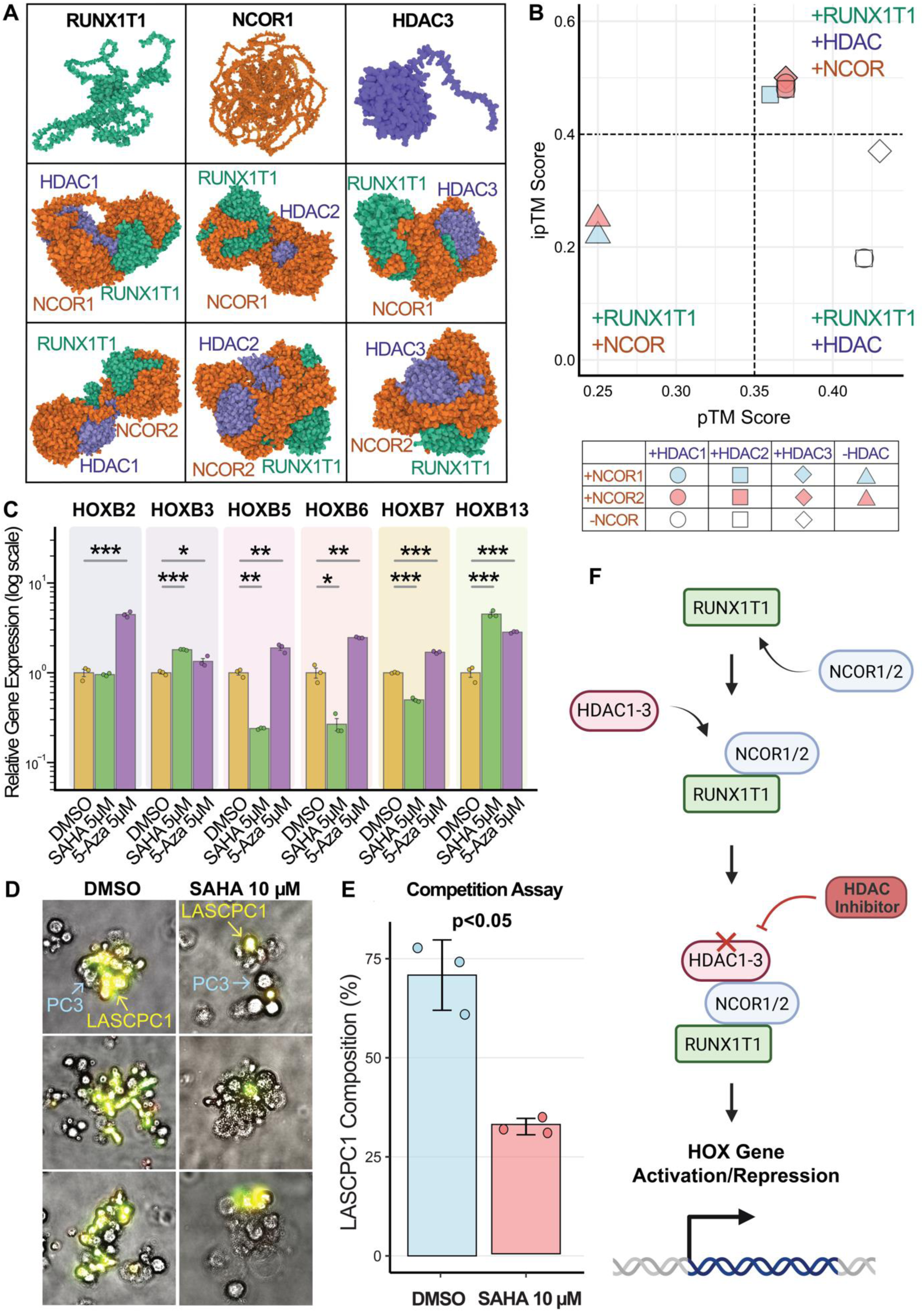
AI-predicted binding partners of RUNX1T1 are NCORs and HDACs, which represent potential therapeutic targets for reversing lineage plasticity. (A) AlphaFold3 AI model prediction showing that RUNX1T1 forms complexes with NCOR1/2 and HDAC1–3. (B) Plot of predicted Template Modeling (pTM) and interface predicted Template Modeling (ipTM) scores for the predicted protein complexes. Higher ipTM scores for the tripartite RUNX1T1–NCOR–HDAC complexes indicate greater confidence in the interaction interface than for the bipartite RUNX1T1–NCOR or RUNX1T1–HDAC complexes. (C) qRT-PCR analysis of HOX gene expression in LASCPC1 cells following 48 h treatment with the HDAC inhibitor Vorinostat (SAHA, 5 µM) or the DNMT inhibitor 5-Azacytidine (5-Aza, 5 µM). SAHA-mediated HDAC inhibition partially reversed HOX code gene expression changes in lineage-plastic cells. (D) Representative microscopy images of a co-culture competition assay. Lineage-plastic LASCPC1 cells labeled with GFP or mCherry fluorescence were co-cultured with lineage non-plastic PC3 cells (unlabeled) and treated with DMSO or SAHA (10 µM) for 7 days. Images were taken on day 2. (E) Quantification of the competition assay by flow cytometry, showing that SAHA-mediated HDAC inhibition significantly reduces the relative proliferation advantage of lineage-plastic LASCPC1 cells compared to non-plastic PC3 cells (p < 0.05). (F) Proposed mechanistic model in which RUNX1T1 recruits the NCOR/HDAC co-repressor complex to chromatin, leading to HOX gene activation or repression, possibly through histone modification. For all panels, significance is denoted as *p < 0.05, **p < 0.01, ***p < 0.001.

Building on these AI-based structural predictions, we next sought functional validation of the RUNX1T1–HDAC interaction. We reasoned that if RUNX1T1 regulates gene expression by recruiting an HDAC-containing corepressor complex, then inhibition of HDAC activity should reverse the RUNX1T1-driven lineage plasticity. To test this, we treated lineage-plastic LASCPC1 cells with the HDAC inhibitor Vorinostat (SAHA) or the DNMT inhibitor 5-Azacytidine (5-Aza) as a negative control, since 5-Aza does not specifically target HDAC. Strikingly, SAHA, but not 5-Aza, restored a non-plastic HOXB code gene expression pattern, recapitulating the siRNA knockdown results and functionally validating the critical role of the HDAC component in this complex (Figure 8C).

Encouraged by these results, we next examined whether this RUNX1T1–HDAC dependency creates a therapeutically actionable vulnerability. We performed a co-culture competition assay using fluorescently labeled lineage-plastic LASCPC1 cells together with non-plastic PC3 cells. While the plastic cells thrived under control conditions (DMSO), SAHA treatment caused a dramatic and selective reduction of their growth in competition with PC3 (Figure 8D, 8E). Taken together, these findings support a mechanistic model (Figure 8F) in which RUNX1T1 recruits the NCOR/HDAC complex to regulate HOX code gene expression, thereby enforcing a lineage-plastic state critically dependent on continuous HDAC activity for survival. This reveals a promising therapeutic strategy to target and suppress plastic, treatment-resistant cancer cells.

## Discussion

Emerging evidence has highlighted the key role of lineage plasticity in driving resistance to many standard cancer therapies across various types of human cancers, including prostate, breast, lung, and pancreatic cancers, as well as melanoma [2, 4-12]. Many known factors driving lineage plasticity have been identified in different cancer subtypes, such as loss of TP53 and RB1 [4, 22], ectopic activation of SOX2, TET2, EZH2, JAK-STAT, and ASCL1 [2, 4, 23, 25, 26, 33], and modification of epigenetic regulators such as CHD1, LSD1, and the SWI/SNF complex [5, 7, 27-32]. Despite these recent advances revealing key regulators of lineage plasticity, a pan-cancer gene marker that quantifies lineage plasticity is still lacking. This absence limits the definition and quantification of lineage plasticity across diverse cancer subtypes and lineages. This knowledge gap significantly restricts the clinical utilization of lineage plasticity as a predictive biomarker for targeted therapy resistance and hinders the practice of precision oncology. Our study demonstrates that the role of the HOX code in representing tissue identity is largely retained in cancer cell lineages. By applying the HOX code to transcriptomic data from over 114 cancer subtypes, we identified lineage-plastic cancers, such as PRSCC, SCLC, and AML, that exhibit distinct HOX codes compared to their non-plastic counterparts. These findings underscore the potential of HOX code analysis in determining the plasticity status of cancer cell lineages, a critical treatment resistance mechanism, which could improve prognosis and inform clinical decisions.

The HOX code, defined by the spatially and temporally coordinated expression of Homeobox (HOX) genes, serves as a critical determinant of embryonic patterning and cell lineage identity during normal development [36]. However, early exploration of the HOX code across pan-cancer subtypes was limited due to insufficient sample sizes. Through comprehensive bioinformatic analysis of multi-omics data, including expression profiles of 39 HOX genes from over 80,000 RNA sequencing samples across 114 cancer subtypes, we demonstrated that HOX codes effectively represent the lineages of cancer cells. Our research represents a significant advancement in translating developmental biology concepts into cancer research applications. While HOX codes have long been recognized for their role in embryonic development and tissue specification, our study expands this concept into cancer biology, providing novel insights into how these developmentally regulated genes contribute to cancer lineage plasticity. By establishing the HOX code as a universal indicator of cancer lineage identity, our work addresses a major limitation in current cancer classification approaches. Unlike canonical cancer lineage markers, which are often loosely defined and vary considerably across different cancer subtypes, HOX code analysis offers a standardized, genome-wide approach to assessing lineage identity regardless of cancer origin. This universality enables more consistent classification of cancer subtypes and more reliable identification of lineage-plastic states across diverse malignancies.

Using this newly developed HOX code, we identified previously uncharted lineage-plastic tumor subtypes in prostate cancer, lung cancer, and AML, which exhibit altered HOX code gene expression patterns compared to their non-plastic counterparts. More notably, our analysis revealed significantly elevated RUNX1T1 expression across all three lineage-plastic cancer subtypes, which was further validated through both bulk and single-cell RNA sequencing data derived from preclinical and clinical samples. Together, our findings provide a novel strategy for characterizing lineage plasticity in pan-cancer cells through HOX code analysis and further suggest ectopic RUNX1T1 expression as a pan-cancer marker and critical mediator of lineage plasticity. We also functionally confirmed the critical but previously uncharted role of RUNX1T1 in lineage plasticity, as siRNA-mediated knockdown of RUNX1T1 reversed key features of the plastic phenotype in multiple preclinical models. Mechanistically, we propose that RUNX1T1 acts by recruiting the NCOR/HDAC co-repressor complex to chromatin, an interaction supported by high-confidence AI-based structural predictions and validated by pharmacological inhibition of HDACs. This addresses a major gap in utilizing lineage plasticity as a predictive biomarker for targeted therapy resistance.

Historically, research on RUNX1T1 has been overwhelmingly focused on its role as part of the RUNX1–RUNX1T1 fusion protein in acute myeloid leukemia. Consequently, the function of the wild-type, unfused RUNX1T1 protein has remained largely understudied, particularly in the context of solid tumors. For the first time, our work reveals a previously unknown and critical role of RUNX1T1 in lineage plasticity, demonstrating that ectopic expression of wild-type RUNX1T1, independent of fusions or mutations, serves as a powerful driver of lineage plasticity across multiple cancer subtypes. This discovery calls renewed attention to a previously overlooked oncogenic driver and highlights its downstream effectors, such as HDACs within its corepressor complex, as novel therapeutic targets for lineage-plastic cancers.

Moreover, our findings have substantial clinical implications, particularly in the context of treatment resistance. Lineage plasticity has emerged as a key mechanism driving resistance to both targeted therapies and immunotherapies. By enabling detection and monitoring of lineage plasticity through HOX code analysis, our study provides a potential strategy for improving patient prognosis and guiding therapeutic decision-making. Clinicians could leverage HOX code signatures to identify patients at risk of developing treatment resistance due to lineage plasticity and adjust treatment regimens accordingly. In addition, our discovery of the RUNX1T1–HDAC axis uncovers a novel therapeutic vulnerability. We demonstrate that lineage-plastic cells are uniquely dependent on this axis, as treatment with an HDAC inhibitor selectively reduces their viability, suggesting a promising and novel strategy to eliminate therapy-resistant populations with potentially minimal side effects.

Collectively, our study establishes HOX code analysis as a powerful approach for detecting cancer lineage plasticity, providing both biological insights into cancer evolution and practical applications in clinical oncology. This work lays a foundation for future investigations of the RUNX1T1–HDAC axis in cancer and supports the clinical evaluation of HDAC inhibitors as a strategy to overcome lineage plasticity–mediated treatment resistance.

## Methods

### Data Sources and Preprocessing

We analyzed bulk RNA sequencing (RNA-seq) datasets from sources including: the pan-cancer cell line transcriptome atlas (PCTA, https://pcatools.shinyapps.io/PCTA_app/), the OHSU AML cohort from cBioportal (https://www.cbioportal.org/), lung cancer patient data from Gene Expression Omnibus (GEO, https://www.ncbi.nlm.nih.gov/geo/). The PCTA dataset comprises TPM (Transcripts Per Million) and read count values for human GRCh38 genes across multiple cancer cell lines. Metadata for PCTA samples, including cell line identifiers, cancer types, and cancer anatomic sites, were retrieved from a previous publication [44]. For single-cell RNA-seq validation, we utilized two prostate cancer datasets: the Human Prostate Single-Cell Atlas (HuPSA) and the Mouse Prostate Single-Cell Atlas (MoPSA) retrieved from previous publication. HUPSA was filtered to exclude normal and normal-adjacent cells, retaining only " PRAD_AR+_1" and "PRSCC" cell types representing the lineage non-plastic and lineage plastic prostate cancer cells, similarly, MOPSA was subset to " PRAD_1" and "PRSCC" cells from the GEM (Genetically Engineered Mouse) model. Both datasets underwent UMAP dimensionality reduction [45]. Additionally, bulk RNA-seq data from the Proatlas provided TPM values for prostate cancer samples, which were subset to RUNX1T1 expression for further analysis [45].

### HOX Code Correlation Analysis

Median HOX gene expression per cancer cell line was calculated, forming a matrix that was hierarchically clustered within cancer type groups using Euclidean distance and Ward.D2 linkage. The resulting correlation matrix was visualized as a heatmap with diagonal preservation using ComplexHeatmap, split by cancer type and annotated with cancer anatomic sites. Similarly, median HOX gene expression per cancer type was calculated and visualized as an upper triangular heatmap.

Pearson correlation coefficients were calculated between HOX gene expression profiles across cancer cell lines and cancer types using the formula:

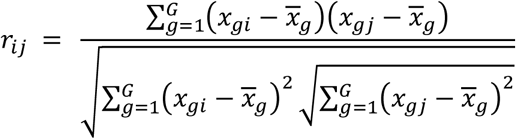

where *x*_*gi*_ is the expression of gene *g* in sample *i*, and 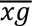 is the average expression of gene *g* across all samples. Prior to computing the correlation matrix, expression values were log-transformed. The distributional properties of the transformed data were assessed using density plots and visual inspection, which confirmed that the use of Pearson correlation was appropriate for measuring pairwise expression similarity.

To further assess lineage-specific HOX code conservation, we grouped correlation values by combined Oncotree lineage and cancer type (e.g., "Neuroendocrine_Prostate"). For each group, Pearson correlations were divided into “Within” and “Between” categories depending on sample pairings, and two-sided Wilcoxon rank-sum tests were applied to compare distributions. Median differences (Within - Between) and p-values were computed across all groups. Groups with *p*<0.05 were labeled significant, and significance was visualized via circular boxplots ordered by effect size. The statistical workflow was implemented in R using the dplyr and ggplot2 packages.

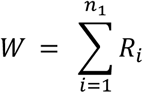

Where *R*_*i*_ is rank of the *i*th value in group 1 after pooling all values and *n*_1_denotes the sample size of group 1. We first calculated Pearson correlation coefficients to measure HOX code similarity between samples. To statistically evaluate whether within-cancer-type correlations were systematically higher than between-type correlations, we applied the one-sided Wilcoxon rank-sum test to the resulting correlation values. Correlation values were not assumed to be normally distributed, and visual inspection confirmed skewed distributions, justifying the use of a non-parametric approach.

### Statistical Detection of Multimodality

To evaluate potential lineage plasticity, we assessed whether the distribution of HOX correlation coefficients within each anatomic origin exhibited multimodality, we applied Hartigan’s Dip Test for unimodality (Hartigan & Hartigan, 1985). The null hypothesis *H*_0_: *f*(*x*) is unimodal was tested against the alternative *H*_1_: *f*(*x*) is multimodal, where Dₙ measures the maximum difference between the empirical distribution function *F*_*n*_(*x*) and the best-fitting unimodal distribution *U*(*x*):

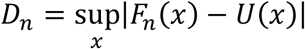

A significant dip statistic (p < 0.05) indicates deviation from unimodality.

In addition, to quantify the number of modes (peaks) in each distribution, we performed local maxima detection on the kernel density estimate *f̂*(*x*), using the Sheather–Jones (SJ) bandwidth selector (Sheather & Jones, 1991) to determine the optimal smoothing parameter *h*_*SJ*_:

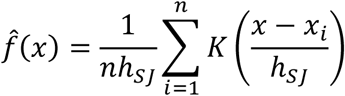

Local maxima of *f̂*(*x*) were counted to estimate the number of distinct modes present in the correlation distribution.

### Differential Expression Analysis

Differential expression analysis was performed using DESeq2 on raw count data from PCTA. Target cell lines were grouped as AML non-plastic (MOLM13, MOLM14), AML plastic (HEL, OCIAML3), prostate non-plastic (22RV1, C42B, PC3), prostate plastic (H660), lung non-plastic (CALU3, H1993), and lung plastic (DMS454, H661, H82). Counts were subset to these groups, prefiltered to retain genes with ≥10 counts in ≥3 samples and analyzed. Results were filtered for upregulated genes using thresholds of log2 fold change (log2FC) > 3 and adjusted p-value (padj) < 0.01, visualized with volcano plots using EnhancedVolcano, highlighting *RUNX1T1*.

DESeq2 models gene-level counts using a negative binomial generalized linear model:

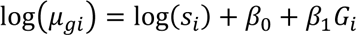

Where *μ*_*gi*_ denotes expect count for gene *g* in sample *i*, *s*_*i*_ is the sample-specific size factor, *G* is the binary indicator of group membership (e.g., plastic vs. non-plastic), and *β*_1_denotes the estimated log2 fold change. We verified the model assumptions by evaluating the dispersion–mean relationship and confirming appropriate normalization via diagnostic plots. These checks confirmed that DESeq2’s negative binomial model was appropriate for the selected count data.

### Statistical Analysis and Visualization

All statistical analyses used R (v4.2.1). Correlation heatmaps employed ComplexHeatmap with colorRamp2 for color gradients. Boxplots and violin plots used ggplot2. We compared RUNX1T1 expression levels across plastic and non-plastic groups using Wilcoxon rank-sum tests, as implemented in stat_compare_means() from ggpubr. Expression values exhibited non-normal distributions based on Q–Q plot assessment, validating the use of non-parametric methods.

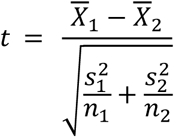

Where *X̄*_1_ and *X̄*_2_ are group means of RUNX1T1 expression, 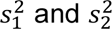 are group variances, and *n*_1_ and *n*_2_ are sample sizes for each group. RUNX1T1 expression values were log2-transformed before group comparisons. Normality was evaluated using Q–Q plots, and equality of variances was tested via Levene’s test. When both assumptions were met, a two-sided Student’s t-test was used. However, if the data showed non-normal distribution or unequal variances, the non-parametric Wilcoxon rank-sum test was used to ensure a robust comparison that is not sensitive to such deviations.

Venn diagrams for overlapping genes were created with VennDiagram. Plots were saved as PDFs with specified dimensions.

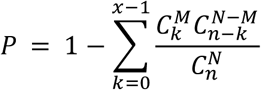

Where *N* denotes total number of genes in the background universe, *M* is the number of genes in a specific pathway, *n* is the number of genes in the input gene list and *x* is the number of overlapping genes between input list and pathway. We used the hypergeometric test to assess statistical enrichment of pathways and gene sets among upregulated genes. The input gene list and background universe were matched for completeness of annotation. Multiple testing corrections were applied using the Benjamini–Hochberg procedure to control false discovery rate (FDR).

### RNA Extraction and Quantitative Real-Time PCR (qRT-PCR)

Total RNA was extracted from cultured cells using Monarch® Total RNA Miniprep Kit (NEB, #T2110) according to the manufacturer’s instructions. RNA concentration and purity were assessed using a NanoDrop spectrophotometer. One microgram of total RNA was reverse transcribed into cDNA using the ABScript III RT Master Mix (Abclonal, RK20429). Quantitative real-time PCR was performed on a QuantStudio 5 System (Applied Biosystems) using the 2X Universal SYBR Green Fast qPCR Mix (ABclonal, RK21203). Gene expression levels were normalized to the housekeeping gene GAPDH. The relative expression was calculated using the comparative Ct (ΔΔCt) method. All experiments were performed with at least three biological replicates. Primer sequences are available upon request.

### Protein Extraction and Western Blotting

Cells were washed with ice-cold PBS and lysed in RIPA buffer (Thermo Fisher Scientific, 89901) supplemented with a benzonase nuclease (Sigma-Aldrich, E1014), protease, and phosphatase inhibitor cocktail (Roche). Protein concentrations were determined using the BCA Protein Assay Kit (Thermo Fisher Scientific). Equal amounts of protein (20-30 µg) were separated by SDS-PAGE on 4-15% Mini-PROTEAN TGX Precast Gels (Bio-Rad) and transferred to a polyvinylidene difluoride (PVDF) membrane (Millipore). Membranes were blocked for 1 hour at room temperature in 5% non-fat dry milk in Tris-buffered saline with 0.1% Tween 20 (TBST). Membranes were then incubated overnight at 4°C with primary antibodies against RUNX1T1 (ABclonal, Cat# A1737, 1:600; Proteintech, Cat# 15494-1-AP, 1:5000) and GAPDH (Cell Signaling Technology, Cat# 14C10, 1:1000) as a loading control. After washing three times with TBST, membranes were incubated with HRP-conjugated secondary antibodies for 1 hour at room temperature. Protein bands were visualized using the SuperSignal West Pico PLUS Chemiluminescent Substrate (Thermo Fisher Scientific, 34577), imaged on a ChemiDoc Imaging System (Bio-Rad), and quantified using Image Lab (Bio-Rad).

### siRNA-Mediated Gene Knockdown

For transient knockdown experiments, a pool of siRNAs targeting human RUNX1T1 and a non-targeting control pool were used (MedChemExpress, HY-RS12350). Cells were seeded in 6-well plates and grown in antibiotic-free medium to 60-80% confluency. On the day of transfection, a 20 µM stock solution of siRNA was prepared. For each well, the siRNA pool was diluted in 100 µL of serum-free DMEM (Solution A). Separately, 6 µL of PolyJet™ In Vitro Transfection Reagent (SignaGen Laboratories) was diluted in 100 µL of serum-free DMEM and immediately added to Solution A. The mixture was incubated for 15 minutes at room temperature to allow for complex formation. The culture medium was removed from the cells and replaced with 900 µL of fresh serum-free DMEM. The 200 µL siRNA-reagent complex was then added dropwise to each well for a final concentration of 50 nM. After an overnight incubation, the medium was replaced with complete culture medium containing serum (RPMI). Cells were harvested for RNA extraction 72 hours post-transfection, and knockdown efficiency was confirmed by qRT-PCR.

### Drug Treatment

For drug inhibition experiments, 1 million LASCPC1 cells were suspended in 3 mL RPMI medium in each well of 6-well plates. DMSO (vehicle control), 5 µM Vorinostat (SAHA, MedChemExpress, HY-10221), or 5 µM 5-Azacytidine (MedChemExpress, HY-10586) were added to 3 biological replicates each. Cells were incubated under 37°C in a CO2 incubator with the inhibitors for 48 hours before being harvested for RNA extraction and subsequent qRT-PCR analysis as described above.

### Co-culture Competition Assay

Fluorescently labeled (GFP, mCherry) LASCPC1 cells were seeded with the non-labeled PC3 cells at a 9:1 ratio (LASCPC1:PC3) in cell culture flasks. The co-cultures were treated with either DMSO (vehicle control) or 10 µM SAHA. Media and drug were refreshed every 2 days. On day 7, cells were collected, washed with PBS, and analyzed on an Attune NxT Acoustic Focusing Cytometer (Invitrogen). Three biological replicates were performed for each condition.

## Code and data availability

All data used in this study is listed in the method section. All code necessary to reproduce every figure in this study is published in GitHub (@JYY2001).

## Acknowledgments

This work was supported or partially supported by: National Cancer Institute/National Institutes of Health: R00CA218885, R37CA258730, R01CA288820, R01CA292949 P. Mu; Department of Defense: W81XWH-18-1-0411 and W81XWH21-1-0520 P. Mu; Cancer Prevention Research Institute (CPRIT): RR170050, RP220473, P. Mu, Prostate Cancer Foundation: 17YOUN12 P. Mu, Yale Cancer Center CCSG Pilot Grant. P30CA016359 P. Mu. and I.Y.K.

## CRediT author statement

**Yuyin Jiang**: Conceptualization, Visualization, Methodology, Investigation, Data Curation, Writing-Original draft preparation; **Siyuan Cheng**: Conceptualization, Methodology, Visualization, Software; **Catherine Yijia Zhang**: Investigation, Data Curation, Visualization; **Xiao Jin**: Formal Analysis, Visualization; **Yaru Xu**: Resources; **Isaac Kim**: Supervision; **Ping Mu**: Conceptualization, Supervision, Writing-Reviewing and Editing.

## Conflict of Interest Statement

P.M. served as a scientific consultant to Accutar Biotechnology, Inc. No other authors have COI to disclose.

During the preparation of this work the author(s) used ChatGPT in order to proofread and correct grammar and typographical errors. After using this tool/service, the author(s) reviewed and edited the content as needed and take(s) full responsibility for the content of the publication.

